# Fibroblast growth factor receptor facilitates recurrence of minimal residual disease following trastuzumab emtansine therapy

**DOI:** 10.1101/731299

**Authors:** Saeed S. Akhand, Stephen Connor Purdy, Zian Liu, Joshua C. Anderson, Christopher D. Willey, Michael K. Wendt

## Abstract

Trastuzumab-emtansine (T-DM1) is an antibody-drug conjugate (ADC) that efficiently delivers a potent microtubule inhibitor into HER2 overexpressing tumor cells. However, resistance to T-DM1 is emerging as a significant clinical problem. Continuous *in vitro* treatment of HER2-transformed mammary epithelial cells with T-DM1 did not elicit spontaneously resistant cells. However, induction of epithelial-mesenchymal transition (EMT) via pretreatment with TGF-β1 facilitated acquisition of T-DM1 resistance. Flow cytometric analyses indicated that induction of EMT decreased trastuzumab binding, prior to overt loss of HER2 expression. Kinome analyses of T-DM1 resistant cells indicated increased phosphorylation of ErbB1, ErbB4, and fibroblast blast growth factor receptor 1 (FGFR1). T-DM1 resistant cells failed to respond to the ErbB kinase inhibitors lapatinib and afatinib, but they acquired sensitivity to FIIN4, a covalent FGFR kinase inhibitor. *In vivo,* T-DM1 treatment led to robust regression of HER2-transformed tumors, but minimal residual disease (MRD) was still detectable via bioluminescent imaging. Upon cessation of the ADC relapse occurred and secondary tumors were resistant to additional rounds of T-DM1, but this recurrent tumor growth could be inhibited by FIIN4. Expression of FGFR1 was upregulated in T-DM1 resistant tumors, and ectopic overexpression of FGFR1 was sufficient to enhance tumor growth, diminish trastuzumab binding, and promote recurrence following T-DM1-induced MRD. Finally, patient-derived xenografts from a HER2^+^ breast cancer patient who had progressed on trastuzumab failed to respond to T-DM1, but tumor growth was significantly inhibited by FIIN4. Overall, our studies strongly support therapeutic combination of TDM1 and FGFR targeted agents in HER2^+^ breast cancer.

## Introduction

Human epidermal growth factor receptor 2 (HER2) is a member of the ErbB family of receptor tyrosine kinases. HER2-amplified breast cancers respond to treatment with the HER2-targeted monoclonal antibodies pertuzumab and trastuzumab at a high rate but acquired resistance to these therapies remains a major clinical problem for patients with this breast cancer subtype. Trastuzumab-emtansine (T-DM1) is an antibody-drug conjugate (ADC) that provides a mechanism to deliver a potent microtubule-targeting cytotoxin to HER2 overexpressing cells. Initial enthusiasm for T-DM1 based on dramatic preclinical results has been diminished by the inability of T-DM1 to improve patient outcomes as compared to the current standard of care therapy, unconjugated trastuzumab in combination with a taxane (1,2). These clinical data suggest that uncharacterized drivers of resistance are at play (3). An underlying mechanism of resistance to all ErbB-targeting compounds is the downregulation of ErbB receptors during the processes of invasion and metastasis. Indeed, clinical findings demonstrate discordance in HER2 expression when metastases are compared to the corresponding primary tumor from the same patient (4,5).

Epithelial-mesenchymal transition (EMT) is a normal physiological process whereby polarized epithelial cells transition into motile, apolar fibroblastoid-like cells to facilitate several developmental events and to promote wound repair in response to damaged tissues (6). In contrast, initiation of pathological EMT engenders the acquisition of invasive, metastatic and drug resistant phenotypes to developing and progressing carcinomas (7)(8,9). Physiologic and pathologic EMT can be induced by cytokines such as TGF-β and HGF (10). More recent findings demonstrate that EMT can be initiated by treatment with kinase inhibitors and that this transition to a mesenchymal state facilitates tumor cell persistence in the sustained presence of these molecular-targeted compounds (11). In contrast to kinase inhibition, very little is known about the mechanisms by which EMT may facilitate resistance to antibody and ADC therapies.

Induction of EMT increases the expression of fibroblast growth factor receptor 1 (FGFR1) (12,13). FGFR1 can also undergo gene amplification and translocation, and elevated expression of FGFR1 is associated with decreased clinical outcomes of breast cancer patients (14)(15,16). Work from our lab and others suggest that upregulation of FGFRs and FGF ligands can serve as resistance mechanisms for tumor cells that were originally sensitive to ErbB and endocrine-targeted therapies (16–20). In addition to enhanced expression of the receptor, our recent studies demonstrate that the processes involved in EMT work en masse to support FGFR signaling through diminution of E-cadherin and enhanced interaction with integrins (21). Several different Type I, ATP-competitive kinase inhibitors against FGFR have been developed, and we and others have demonstrated their *in vivo* efficacy in delaying the growth of metastatic breast cancers (12)(22,23). Based on the potential of FGFR as a clinical target for cancer therapy we recently developed FIIN4, a highly specific and extremely potent covalent kinase inhibitor of FGFR, capable of *in vivo* tumor inhibition upon oral administration in rodent models (20,24).

In the current study, we address the hypothesis that FGFR can act as a driver of resistance to T-DM1. The use of *in vivo* and *in vitro* models demonstrate that unlike ErbB-targeted kinase inhibitors, EMT cannot overtly elicit resistance to T-DM1. Instead, induction of EMT and upregulation of FGFR1 induce a cell population with reduced trastuzumab binding. This minimal residual disease (MRD) is able to persist in the presence of T-DM1 and eventually reemerge as recurrent tumors that lack HER2 expression. In this recurrent setting, FGFR acts as a major driver of tumor growth, which can be effectively targeted with FIIN4. Overall, our studies strongly suggested that combined therapeutics targeting HER2 and FGFR will delay tumor recurrence and prolong response times of patients with HER2^+^ breast cancer.

## Methods

### Cell culture and reagents

Bioluminescent, HER2-transformed HMLE cells (HME2) were constructed as previously described (20). HME2 and BT474 cells stably overexpressing FGFR1 were also previously described (20). Trastuzumab and trastuzumab emtansine (T-DM1) were obtained from Genentech Biotechnology Company through the material transfer agreement program. Where indicated HME2 cells were treated with TGF-β1 (5 ng/ml) every three days for a period of 4 weeks to induce EMT. These EMT-induced HME2 cells were further treated with T-DM1 (250 ng/ml) every three days until resistant colonies emerged, these cells were pooled and cultured as the TDM1R population. Cells were validated for lack of mycoplasma contamination using the IDEXX Impact III testing on July 24th, 2018.

### Xenograft studies and drug treatments

HME2 cells (2×10^6^) expressing firefly luciferase were injected into the duct of the 4^th^ mammary fat pad of female NRG mice. When tumors reached a size of 200 mm^3^, mice were treated with the indicated concentrations of T-DM1 via tail vein injection. Presence of tumor tissue was visualized by bioluminescence imaging following I.P. administration of luciferin (Gold Bio). Where indicated viable pieces of HME2 tumor tissue were directly transplanted into the exposed fat pad of recipient NRG mice. Similarly, pieces of human derived HCI-012 PDX (Huntsman Preclinical Research Resource) were engrafted onto the exposed mammary fat pad of female NRG mice. Tumor bearing mice were treated with T-DM1 as indicated followed by FIIN4 (25 mg/kg/q.o.d) resuspended in DMSO and then further diluted in a solution 0.5% carboxymethyl cellulose and 0.25% Tween-80 to a final concentration of 10% DMSO, for administration to animals via oral gavage. Mammary tumor sizes were measured using digital calipers and the following equation was used to approximate tumor volume (V=(length^2^)*(width)*(0.5)). All animal experiments were conducted under IACUC approval from Purdue University.

### Cell Biological assays

For *ex-vivo* 3D culture, viable human PDX tissues were dissected as above but instead of transfer onto recipient animals, pieces were further mechanically dissected and treated with trypsin-EDTA. These cells were shaken several times and incubated at 37° C. Cells were then filtered through a 50 μM filter and plated onto a 50 μl bed of growth factor reduced cultrex (Trevigen) in a white walled 96 well plate. These cultures were allowed to grow for 20 days in the presence or absence of the indicated kinase inhibitors (1 μM). Two-dimensional cell growth dose response assays were conducted in white walled 96 well plates. Cells (5000 cell/well) were plated in the presence of the indicated concentrations of T-DM1 or kinase inhibitors and cultured for 96 hours. In both cases cell viability was determined by Cell Titer Glo assay (Promega).

### Immuno-assays

For immunoblot assays, lysates were generated using a modified RIPA buffer containing 50mM Tris pH 7.4, 150 mM NaCl, 0.25% Sodium deoxycholate, 0.1% SDS, 1.0% NP-40, containing protease inhibitor cocktail (Sigma), 10 mM Sodium Orthovanadate, 40 mM β-glycerolphosphate, and 20 mM Sodium Fluoride. Following SDS PAGE and transfer, PVDF membranes were probed with antibodies specific for FGFR1 (9740), HER2 (4290; Cell signaling technologies), E-cadherin (610182), N-cadherin (610920), Vimentin (550513; BD biosciences), or β-Tubulin (E7-s; Developmental Studies Hybridoma Bank). For immunocytochemistry, formalin fixed paraffin embedded tissue sections were deparaffinized and stained with antibodies specific for HER2 (4290), FGFR1 (HPA056402; Sigma), Ki67 (550609; BD biosciences) or were processed using the TUNEL Assay Kit (ab206386). Additionally, cells were trypsinized and incubated with antibodies specific for FITC conjugated CD44 (338804) and PerCP conjugated CD24 (311113; Biolegend) or trastuzumab conjugated with Alexafluor 647 according to the manufacturer’s instructions (A20181; Thermo Scientific). Following antibody staining these cells were washed and analyzed by flow cytometry.

### Kinomic analyses

Lysates from HME2 cell conditions indicated above, were analyzed on tyrosine chip (PTK) and serine-threonine chip (STK) arrays using 15ug (PTK) or 2ug (STK) of input material as per standard protocol in the UAB Kinome Core as previously described (25,26) Three replicates of chip-paired samples were used and phosphorylation data was collected over multiple computer controlled kinetic pumping cycles, and exposure times (0,10,20,50,100,200ms) for each of the phosphorylatable substrates. Slopes of exposure values were calculated, log2 transformed and used for comparison. Raw image analysis was conducted using Evolve2, with comparative analysis done in BioNavigator v6.2 (PamGene, The Netherlands).

### Statistical analyses

Data from the Long-HER study were obtained from GSE44272. Expression values of FGFR1-4 were obtained from Affymetrix probes, 11747417_x_at, 11740159_x_at, 11717969_a_at, 11762799_a_at, respectively. Expression values were normalized to the average probe value for the entire group and differences between the long term responders (Long-HER) and control (Poor response) groups were compared via an 2-sided, unpaired T-test. 2-way ANOVA or 2-sided T-tests were used where the data met the assumptions of these tests and the variance was similar between the two groups being compared. No exclusion criteria were utilized in these studies. A Log-rank test was performed to calculate statistically significant differences in disease-free survival of HME2-GFP and HME2-FGFR1 tumor–bearing mice. *P* values for all experiments are indicated, values of less than 0.05 were considered significant.

## Results

### HER2 is diminished following recurrence of T-DM1 resistant minimal residual disease

Human mammary epithelial (HMLE) cells can be transformed by overexpression of wild type HER2 (20,27). Engraftment of these cells onto the mammary fat pad results in robust formation of highly differentiated, nonmetastatic, secretory tumors that form a liquid core (26). We engrafted the HER2 transformed HMLE cells (HME2) onto the mammary fat pad and upon formation of orthotopic tumors mice received four intravenous injections of T-DM1 administered once a week for four weeks (Fig. 1A,1B). This treatment protocol led to robust regression of these tumors to a point which they were no longer palpable and therefore immeasurable by digital calipers (Fig. 1A). However, these HME2 cells were constructed to stably express firefly luciferase and MRD was still detectable via bioluminescent imaging (Fig. 1B). Cessation of T-DM1 treatment led to recurrence of mammary fat pad tumors in 3 of 5 mice over approximately a 150-day period (Fig. 1A). Importantly, these recurrent tumors were nonresponsive to additional rounds of T-DM1 (Fig. 1A). Histological assessment of the recurrent tumors clearly demonstrated reduced levels of HER2 as compared to the untreated HME2 tumors (Fig. 1C). Overall these data demonstrate that even cells specifically transformed by HER2 overexpression are capable of establishing drug persistent MRD in response to T-DM1 and recurring in a HER2-independent manner.

**Figure 1.**
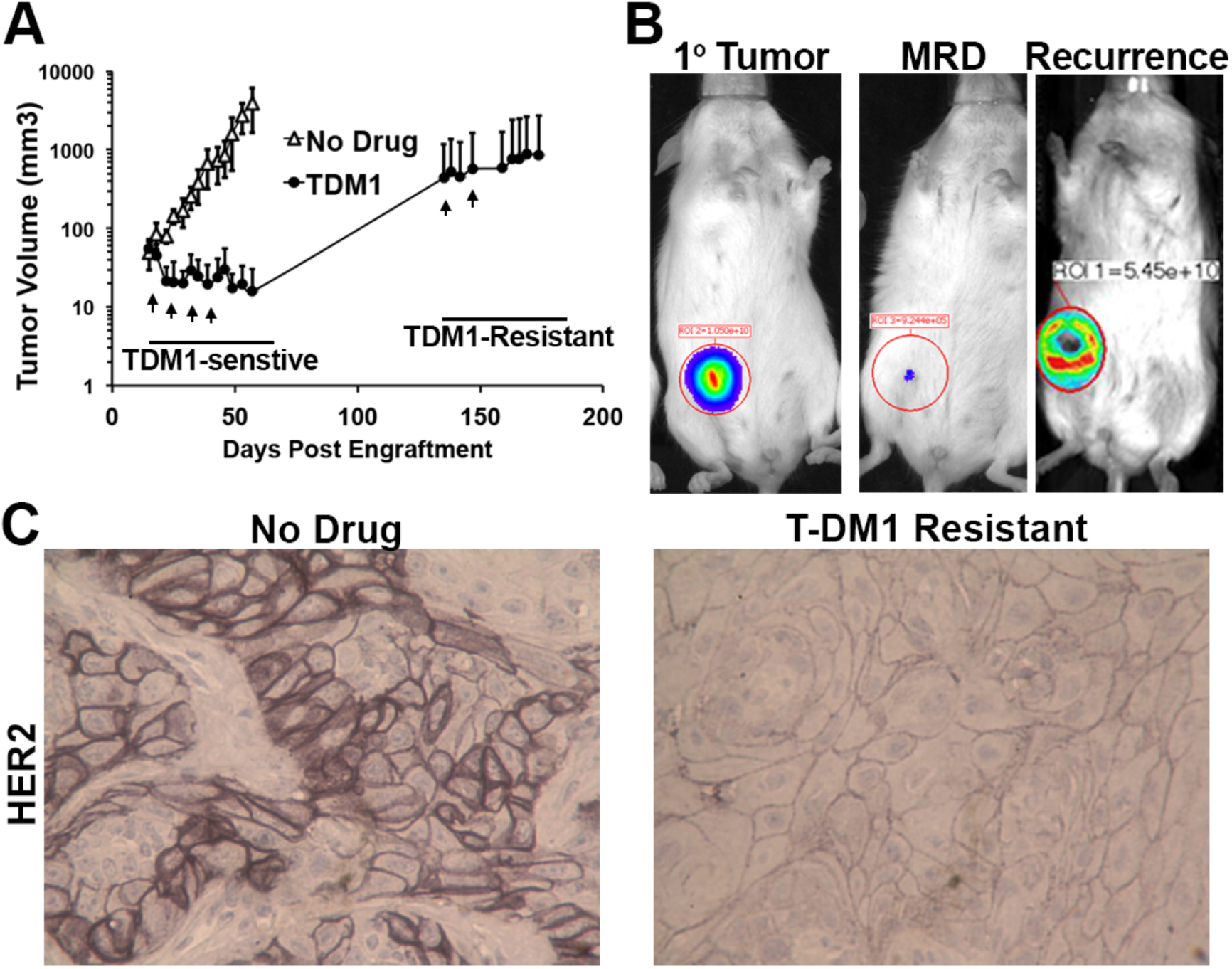
HER2 is diminished following recurrence of T-DM1 resistant minimal residual disease. **A,** Mice were inoculated with HME2 cells (2×10^6^ cells/mouse) via the mammary fat pad and tumors were allowed to form for a period of 14 days. At this point mice were split into two cohorts (5 mice/group) and left untreated (no drug) or were treated with T-DM1 (9 mg/kg) at the indicated time points (arrows). Following complete regression of palpable tumors T-DM1 treatment was stopped. Recurrent tumors (3 mice) were again treated with T-DM1 (arrows). **B,** Representative bioluminescent images of tumor bearing mice before T-DM1 treatment (1° tumor), following T-DM1 treatment (MRD), and upon tumor recurrence. **C,** Immunohistochemistry for HER2 expression in control and recurrent, T-DM1 resistant HME2 tumors.

### In vitro establishment of T-DM1 resistant cells requires prior induction of epithelial-mesenchymal transition

Attempts to subculture the T-DM1 recurrent HME2 tumors were unsuccessful, suggesting that these cells had evolved mechanisms of tumor growth that were not present under *in vitro* culture conditions (Suppl. Fig. S1A). Therefore, we sought to establish a T-DM1 resistant cell line via prolonged *in vitro* ADC treatment. However, progressive treatment of HME2 cells with T-DM1 over extended periods of time failed to yield a spontaneously resistant population (Fig. 2A). We recently demonstrated that induction of EMT in the HME2 model is sufficient to facilitate immediate resistance to the ErbB kinase inhibitors, lapatinib and afatinib (20). In contrast, induction of EMT via pretreatment with TGF-β1 did not induce immediate resistance to T-DM1 (Fig 2A; 2 weeks). Consistent with the inhibition of microtubules being the mechanism of emtansine, treatment of parental and TGF-β1 pretreated HME2 cells with T-DM1 prevented cell division leading to the formation of non-dividing groups of cells and large senescent cells (Fig. 2A, 2B). Importantly, only those cells that had undergone EMT via pretreatment with TGF-β were capable of giving rise to extremely mesenchymal daughter cells that were capable of replicating in the continued presence of T-DM1 (Fig. 2A, 2B). This *in vitro*-derived T-DM1 resistant (TDM1R) cell population continued to thrive in culture and maintained their mesenchymal phenotype and resistance to T-DM1 even after several passages in the absence of the drug (Fig. 2C). To gain insight into the mechanisms by which induction of EMT facilitates acquisition of resistance to T-DM1 we fluorescently labeled trastuzumab and utilized flow cytometry to quantify changes in drug binding. Induction of EMT with TGF-β clearly produced a distinct population of cells that were resistant to trastuzumab binding, giving rise to a more uniform reduction in trastuzumab binding in the TDM1R cell population (Fig. 2D). These findings suggest that prior induction of cytokine-mediated EMT contributes to loss of trastuzumab binding and is required for acquisition of resistance to T-DM1.

**Figure 2.**
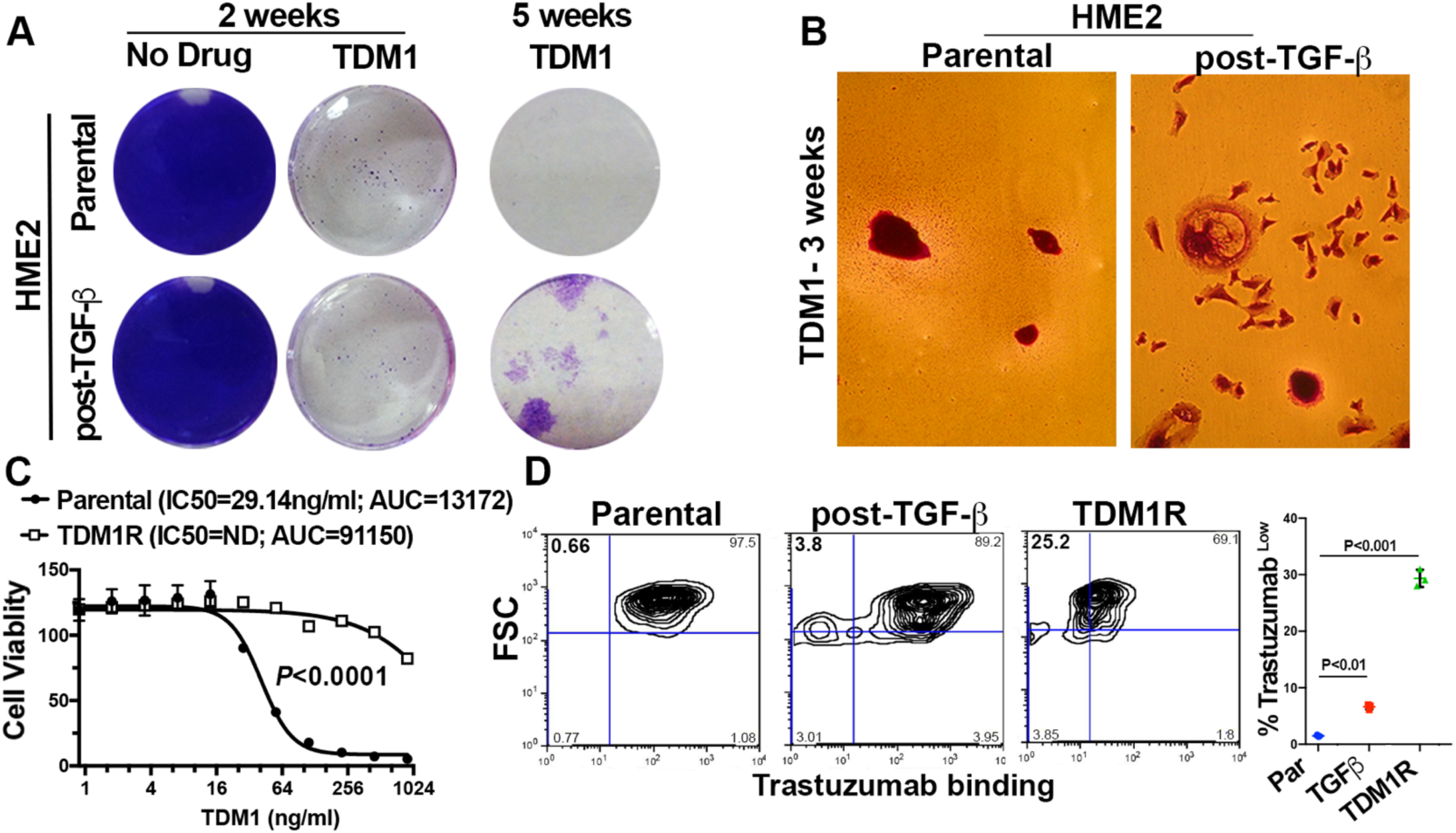
*In vitro* establishment of T-DM1 resistant cells requires prior induction of epithelial-mesenchymal transition. **A,** HME2 cells were left untreated (parental) or were stimulated with TGF-β1 and allowed to recover (Post-TGF-β) as described in the materials and methods. These two cell populations were subsequently treated with T-DM1 (250 ng/ml) every three days for a period of 5 weeks. Representative wells were stained with crystal violet at the indicated time points to visualize viable cells. **B,** Brightfield microscopy of crystal violet stained HME2 parental and post-TGF-β cells following 3 weeks of continuous T-DM1 treatment. **C,** The T-DM1 resistant (TDM1R) cells that survived 5 weeks of treatment were further expanded and cultured for a period of 4 weeks in the absence of T-DM1. These cells along with passage-matched parental HME2 cells were subjected 96 hour treatments with the indicated concentrations of T-DM1 and assayed for cell viability. Data are the mean ±SE of three independent experiments resulting in the indicated P value. **D,** Parental, post-TGF-β, and TDM1R HME2 cells were stained with Alexafluor 647-labeled trastuzumab and antibody binding was quantified by flow cytometry. The percentage of cells in each quadrant with reference to forward scatter (FSC) is indicated. Also shown is the mean, ±SD, percentages of low trastuzumab binding (Trastuzumab^Low^) cells of three independent experiments resulting in the indicated P values.

### FGFR1 is sufficient to reduce T-DM1 binding and efficacy

Our previous studies establish that following TGF-β1 treatment, the purely mesenchymal HME2 culture will asynchronously recover producing a heterogenous population of both epithelial and mesenchymal cells (20). These morphologically distinct populations can also be readily visualized via flow cytometric analyses for CD44 and CD24 (Fig. 3A and 3B). Consistent with the stable mesenchymal morphology of the TDM1R cells they presented as a single population with high levels of CD44, but lacked the diminished expression of CD24 characteristic of TGF-β-induced EMT (Fig. 3A and 3B)(27). In contrast, other markers of EMT were enhanced upon acquisition of T-DM1 resistance, including loss of E-cadherin and potentiated gains in N-cadherin and vimentin (Fig. 3C). Consistent with the diminished trastuzumab binding observed in figure 2, we also observed HER2 expression to be decreased in whole cell lysates from TDM1R cells (Fig. 3C). To elucidate a mechanistic characterization of potential mediators of T-DM1 resistance we compared the TDM1R cells to their T-DM1 sensitive, post-TGFβ HME2, counterparts using kinomic profiling on the PamStation-12 platform. Lysates from TDM1R cells had an increased ability to phosphorylate peptides from FKBP12-rapamycin associated protein (FRAP), a result consistent with enhanced PI3 kinase-mTOR signaling (Suppl. Table 1). This finding is supported by the Long-HER study, which compared global gene expression of HER2^+^ patients that experienced a long-term response to trastuzumab with those whose disease progressed within the first year of initiating trastuzumab (28). Looking upstream to receptors potentially responsible for these events, we observed the autophosphorylation sites of several other ErbB receptors, VEGFRs and FGFRs to be increased in the TDM1R lysates (Suppl. Table 1). Upon further investigation into the potential of these receptors in facilitating resistance to T-DM1 we found that the expression levels of FGFR1 induced by TGF-β were further enhanced upon acquisition of T-DM1 resistance (Fig. 3C). Directed analysis of the Long-HER dataset indicated that enhanced expression of FGFR1 and FGFR2 are significantly associated with a poor clinical response to trastuzumab (Fig. 3D). Along these lines analysis of GSE95414 comparing T-DM1 sensitive NCI-N87 cells to their T-DM1 resistant counterparts indicated potential gains in the expression of FGFR2, 3 and 4 (Suppl. Fig. 2a). In contrast, analysis of GSE100192 indicated that T-DM1 resistant clones derived from the HER2 amplified BT747 cancer cell line do not demonstrate increases in any of the FGFRs (Suppl. Fig. 2B). These data are consistent with recent findings that indicate the BT474 model amplifies the genomic cluster of FGF ligands 3/4/19 instead of modulating FGF receptors (18). To elucidate if FGFR1 is sufficient to provide resistance to T-DM1 we constructed HME2 and BT474 cells to specifically overexpress FGFR1 in the absence of other EMT-associated factors (Fig. 3C; (20)). Using this approach we found that overexpression of FGFR1, when in the presence of exogenous ligand, was sufficient to significantly reduce the dose response to T-DM1 (Fig. 3E; Suppl. Fig. S3). Furthermore, FGFR1 overexpression, in the absence of other EMT-associated events, was sufficient to cause a significant reduction in trastuzumab binding (Fig. 3F).

**Figure 3.**
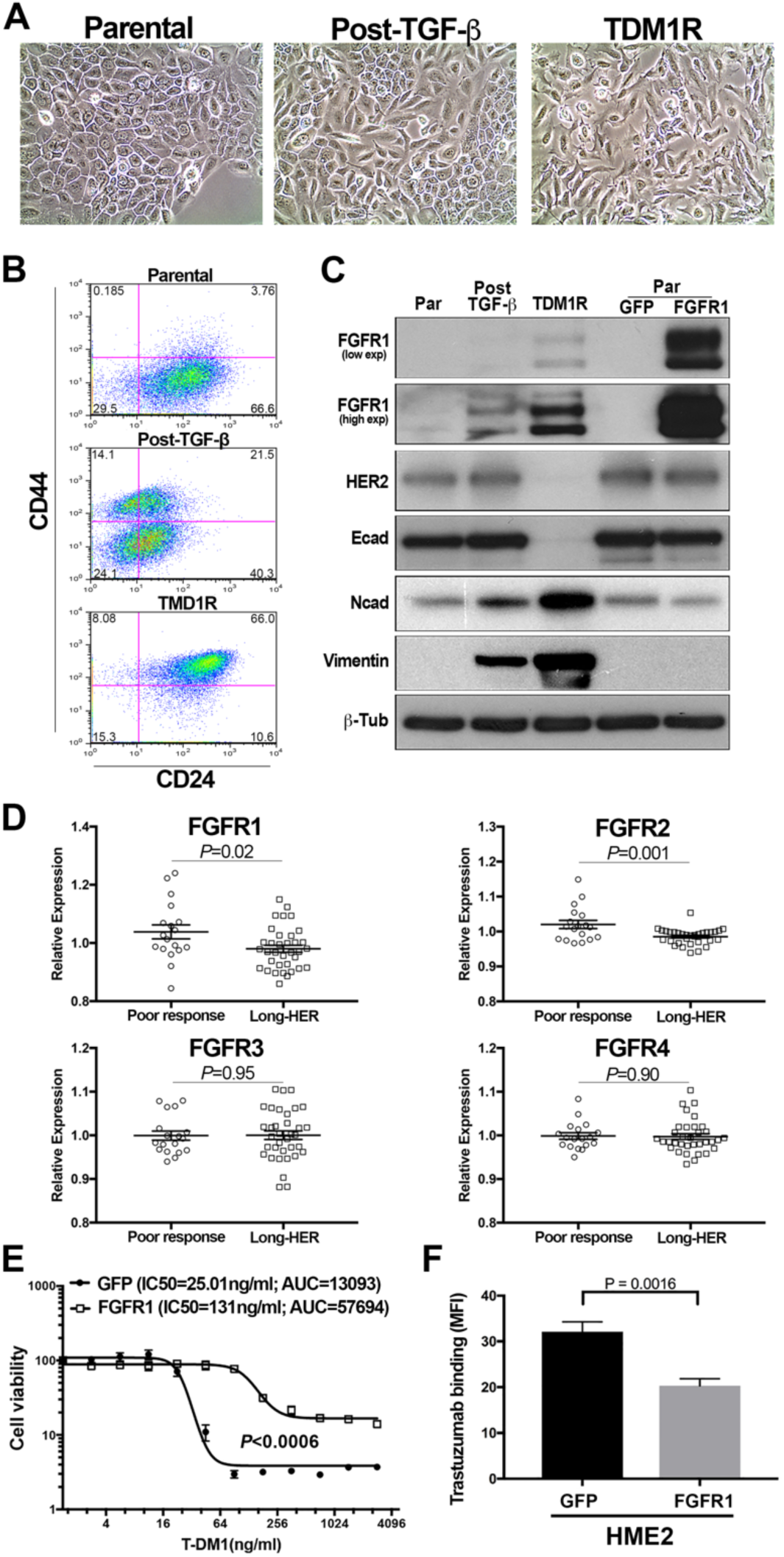
FGFR1 is sufficient to reduce T-DM1 efficacy. **A,** Brightfield microscopy of HME2 parental, post-TGF-β, and T-DM1 resistant (TDM1R) cells. **B,** The cells described in panel A were analyzed by flow cytometry for cell surface expression of CD24 and CD44. The percentage of cells in each quadrant is indicated. **C,** Whole cell lysates from HME2 parental, Post-TGF-β, T-DM1 resistant (TDM1R), and HME2 cells constructed to stably express FGFR1 or GFP as a control were analyzed by immunoblot for expression of FGFR1, HER2, E-cadherin (Ecad), N-cadherin (Ncad), vimentin, and β-tubulin (β-Tub) served as a loading control. Data in panels B and C are representative of at least three separate analyses. **D,** Expression values for FGFR1-4 were analyzed in the Long-HER data set. Data are the relative expression of individual patients that demonstrated long-term (Long-HER) or short-term (Poor-response) response to trastuzumab treatment, resulting in the indicated *P* values. **E,** HME2 cells expressing FGFR1 or GFP as a control were treated with the indicated concentrations of T-DM1 for 96 hours at which point cell viability was quantified. Data are normalized to the untreated control cells and are the mean ±SE of three independent experiments resulting in the indicated *P* value. **F,** HME2 cells expressing FGFR1 or GFP as a control were incubated with Alexafluor 647-labeled trastuzumab and antibody binding was quantified by flow cytometry. Data are the mean fluorescence intensities ±SD for three independent experiments resulting in the indicated *P* value.

### FGFR1 increases tumor recurrence following T-DM1-induced minimal residual disease

We next sought to evaluate the impact of FGFR1 expression on HME2 tumor growth and response to T-DM1. Overexpression of FGFR1 promoted a significant increase in growth rate of HME2 tumors upon mammary fat pad engraftment, leading to differential TDM1 treatment initiation times for matched tumor sizes (Fig. 4A). Irrespective of FGFR1 expression, the liquid filled masses characteristic of large HME2 tumors quickly became necrotic after a single dose of TDM1 (Fig 4B). However, following this initial rapid reduction in tumor size, only the FGFR1 overexpressing tumors maintained a more solid mass which required two additional rounds of T-DM1 treatment to achieve complete tumor regression (Fig. 4B). As we observed in Figure 1 the MRD associated with these nonpalpable lesions could still be detected by bioluminescence (Fig. 4B). Following achievement of T-DM1-induced MRD, none of the control tumors progressed within the 40 day post-treatment observation period (Fig. 4C). In contrast, over 50% of the FGFR1 overexpressing tumors underwent disease progression during this same post-treatment time frame (Fig. 4C). Taken together, these data strongly suggest that enhanced expression of FGFR1 inhibits T-DM1 binding, facilitating therapeutic resistance and post-treatment tumor recurrence.

**Figure 4.**
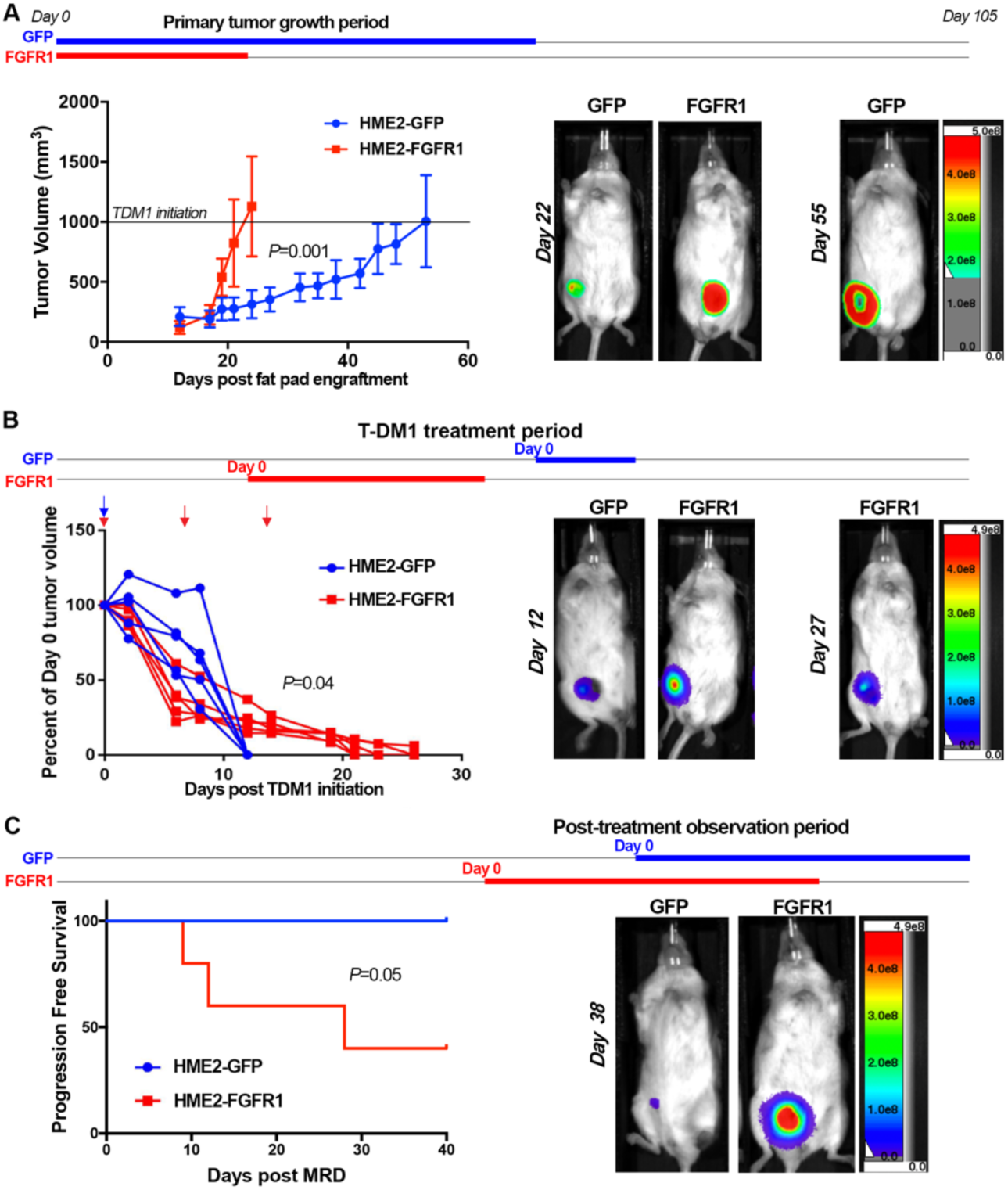
FGFR1 increases tumor recurrence following T-DM1-induced minimal residual disease **A,** Time line demarking the primary tumor growth period for HME2 cells constructed to express FGFR1 or GFP as a control. Cells were engrafted onto the mammary fat pad of female NRG mice (2×10^6^ cells/mouse; n=5 mice per group). Tumor growth was monitored via digital caliper measurements at the indicated time points. T-DM1 treatment was initiated in each group when tumors reached an average of 1000 mm^3^, horizontal line. Representative bioluminescent images for each group are shown at the indicated time points. Days are in reference to tumor engraftment (Day 0). **B,** Time line demarking the T-DM1 treatment periods for each group. T-DM1 was administered via I.V. injections (9 mg/kg) at day 0 for HME2-GFP tumors and days 0,7 and 14 for HME2-FGFR1 tumors (arrows). Tumor regression was monitored via digital caliper measurements at the indicated time points. Representative bioluminescent images for each group are shown at the indicated time points. Days are in reference to initiation of T-DM1 treatment (Day 0) for each group. **C,** Time line demarking the post-treatment observation period for each group. Once tumors regressed to a non-palpable state of minimal residual disease (MRD) mice were left untreated and monitored for tumor recurrence via digital caliper measurements. Representative bioluminescent images for each group are shown at the indicated time points. Days are in reference to achievement of MRD (Day 0) for each group. Data in panel C were analyzed via a log rank test where tumor recurrence of >50 mm^3^ was set as a criteria for disease progression. Data in panel A are the mean ±SD of five mice per group resulting in the indicated P value. In panel B tumor size for each mouse is plotted individually. Data in panels A and B were analyzed via a two-way ANOVA.

### T-DM1 resistant cells are sensitive to covalent inhibition of FGFR

Given the changes in ErbB kinase signaling observed in the TDM1R cells and the ability of FGFR overexpression to facilitate recurrence following ADC therapy we next sought to evaluate the ability of specific kinase inhibitors to target TDM1R cells as compared to their T-DM1 sensitive counterparts. Lapatinib is a clinically used kinase inhibitor that targets both EGFR and HER2, and we recently developed FIIN4, a covalent kinase inhibitor that targets FGFR1-4 (24)(29). Treatment of the HME2 parental cells with lapatinib led a robust inhibition of HER2 phosphorylation and downstream blockade of ERK1/2 phosphorylation (Fig 5A). Consistent with the reduced expression of total HER2, phosphorylated levels of HER2 were undetectable in the TDM1R cells and ERK1/2 phosphorylation was minimally inhibited by lapatinib (Fig 5A). In contrast, treatment of the HME2 parental cells with FIIN4 had no effect on HER2 or ERK1/2 phosphorylation, but FIIN4 markedly diminished ERK1/2 phosphorylation in the TDM1R cells (Fig 5A). Importantly, TDM1R cells also demonstrated robust resistance to lapatinib, even thought they had never been exposed to this compound previously (Fig. 5B). Similarly, TDM1R cells were also resistant to afatinib, a more potent second-generation covalent kinase inhibitor capable of targeting EGFR, HER2, and ErbB4 (Fig. 5C;(30)). In contrast, HME2 cells that had previously been selected for resistance to lapatinib (LAPR) maintained expression of HER2 and were similarly sensitive to T-DM1 as compared to the HME2 parental cells (Fig. 5D;(20)). Finally, TDM1R cells were significantly more sensitive to FIIN4 as compared to the HME2 parental cells (Fig. 5E). Taken together these data indicate that resistance to HER2-targeted ADC therapy predicates acquisition of resistance to ErbB-targeted kinase inhibitors but the reverse is not true. Importantly, these drug resistant populations become increasingly sensitive to covalent inhibition of FGFR.

**Figure 5.**
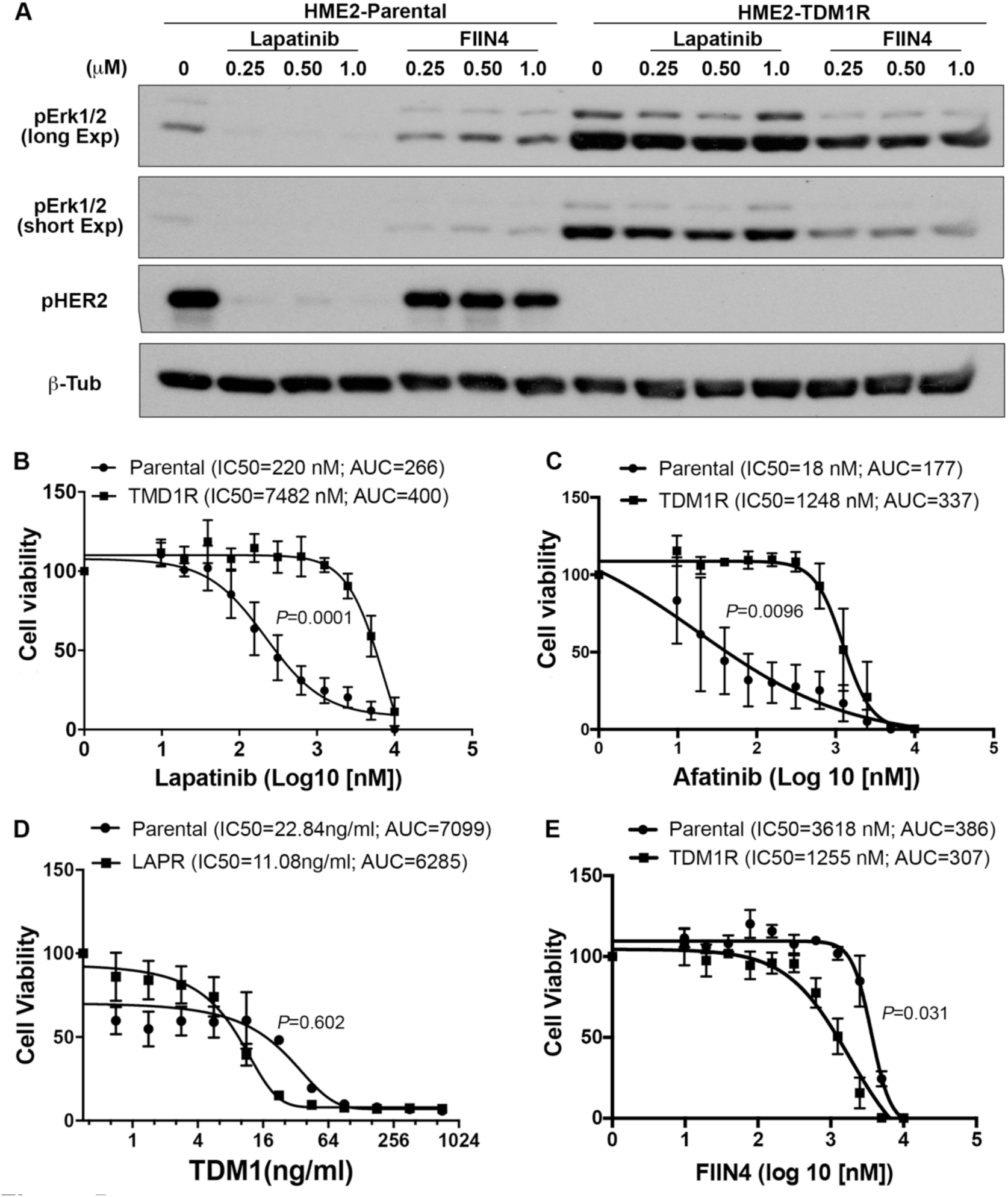
T-DM1 resistant cells are sensitive to covalent inhibition of FGFR. **A,** HME2 parental and T-DM1 resistant (TDM1R) cells were treated with the indicated concentrations of lapatinib or FIIN4 for 2 hours. Cell lysates were subsequently assayed by immunoblot for phosphorylation of ERK1/2 and HER2, β-tubulin (β-Tub) served as a loading control. Data in panel A are representative of at least two independent experiments. **B,** HME2 parental cells (parental) and TDM1R cells were plated in the presence of the indicated concentrations of lapatinib for 96 hours at which point cell viability was determined. **C,** HME2 parental and TDM1R cells were plated in the presence of the indicated concentrations of afatinib for 96 hours at which point cell viability was determined. **D,** HME2 parental and lapatinib resistant (LAPR) cells were plated in the presence of the indicated concentrations of T-DM1 for 96 hours at which point cell viability was determined. **E,** HME2 parental and TDM1R cells were plated in the presence of the indicated concentrations of FIIN4 for 96 hours at which point cell viability was determined. Data in panels B-E are the mean ±SE of at least three independent experiments resulting in the indicated P values.

### T-DM1 resistant tumors respond to systemic inhibition of FGFR

We next sought to validate our *in vitro* findings by evaluating the efficacy of FGFR inhibition in the treatment of tumors that had acquired *in vivo* resistance to T-DM1. To do this we treated HME2 tumor bearing mice with T-DM1 and passaged sections of the remaining tumors onto additional mice. The process was repeated twice until we obtained growing tumors that did not respond to T-DM1 (Fig. 6A). Similar to what was observed in tumors that were recovered following induction of MRD and in our *in vitro* TDM1R cells, these serially passaged *in vivo*-derived T-DM1 resistant tumors also demonstrated a diminution in HER2 expression (Suppl. Fig. 1C). Furthermore, following three rounds of treatment with T-DM1 the resultant tumors expressed readily detectable levels FGFR1 as compared to untreated HME2 tumors (Fig. 6A). Importantly, we were able to significantly inhibit the growth of these T-DM1-resistant tumors upon treatment with FIIN4 (Fig. 6B-6D). This inhibition of tumor growth was consistent with induction of apoptosis and decreased proliferation as visualized by TUNEL and Ki67 staining in tumors from FIIN4 treated mice as compared to untreated controls (Fig. 6E).

**Figure 6.**
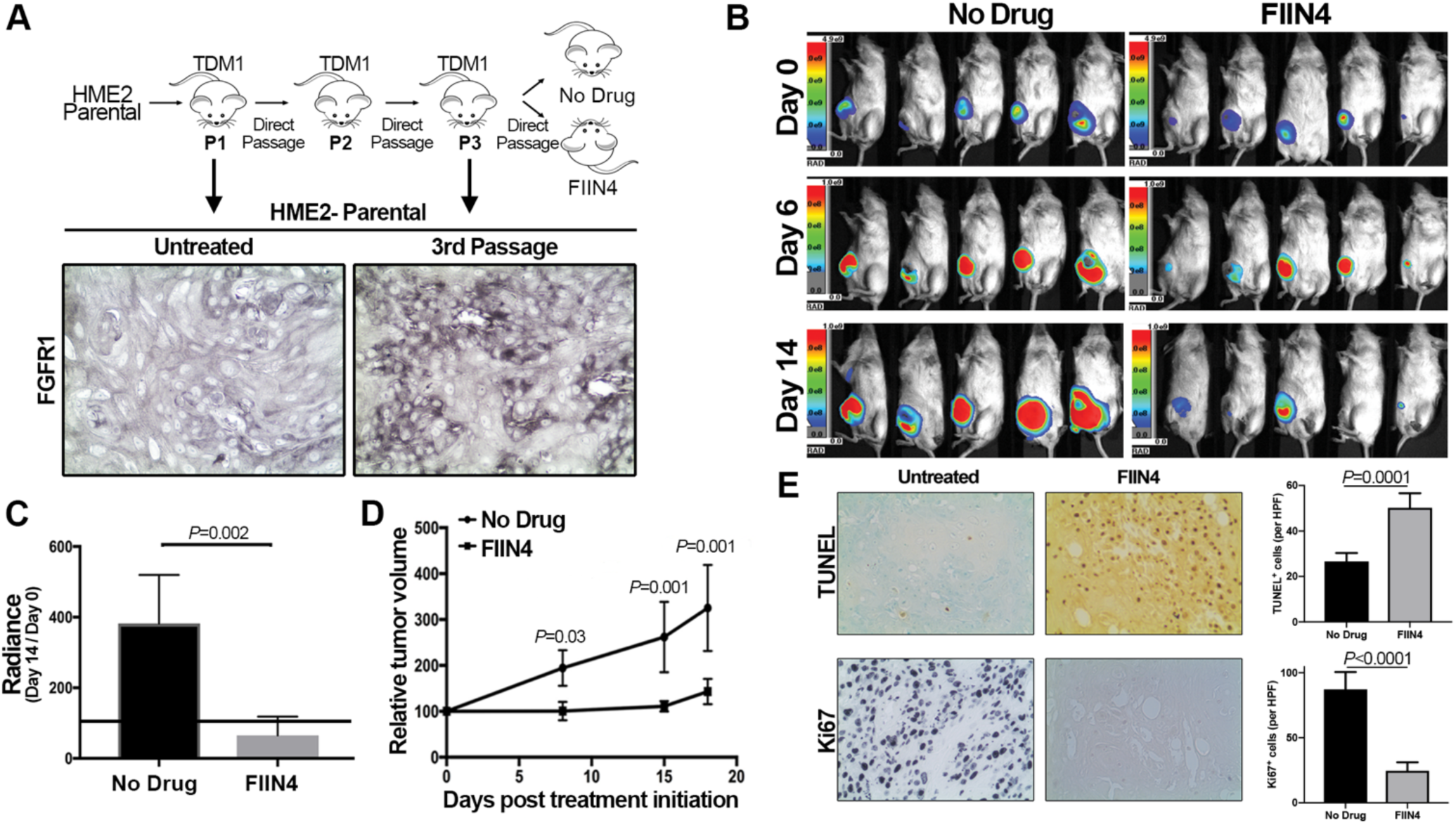
T-DM1 resistant tumors respond to systemic inhibition of FGFR. **A,** Schematic representation of the *in vivo* derivation of T-DM1 resistant HME2 tumors. HME2 parental cells (2×10^6^) were engrafted onto the mammary fat pad of an NSG mouse. This mouse was treated with T-DM1 until tumor regression was observed. Sections of the remaining tumor were directly transferred onto recipient mice. This was repeated twice until transferred tumors that no longer responded to T-DM1 therapy were identified. At this point sections of a T-DM1 resistant tumor were assessed by IHC for FGFR1 expression as compared to the originally engraphed HME2 tumors. Representative FGFR1 IHC staining is shown. **B,** Bioluminescent imaging of mice bearing the T-DM1 resistant tumors described in panel A. These mice were left untreated (No Drug) or were treated with FIIN4 (100 mg/kg/q.o.d.). **C,** Bioluminescent quantification of control and FIIN4-treated animals bearing T-DM1 resistant tumors. Data are normalized to the tumor luminescence values at the initiation of FIIN4 treatment (day 0). **D,** Tumor size as determined by digital caliper measurements at the indicated time points during FIIN4 treatment. For panels C and D data are the mean ±SE of five mice per group resulting in the indicated P-values. **E,** Representative Ki67 and TUNEL staining of P3, T-DM1 resistant tumors from untreated and FIIN4 treated groups. Also shown are the mean ±SD of TUNEL and Ki67 positive cells per high powered field (HPF), n=5, resulting in the indicated P values.

Next, we utilized a patient-derived xenograft (PDX) that was isolated from a pleural effusion of a patient originally diagnosed with a HER2^+^ primary tumor that went on to fail on trastuzumab therapy (Fig. 7A). Consistent with these clinical data and our findings, these PDX tumors displayed minimal staining for HER2 and mice bearing these PDX tumors failed to respond to T-DM1 treatment (Fig. 7B, 7C). These PDX tumors also demonstrated readily detectable staining for FGFR1 (Fig. 7C). Clinically, lapatinib is indicated as a second line therapy in HER2^+^ patients that do not respond to trastuzumab. Treatment with lapatinib did blunt the 3D invasive phenotype of the HCI-012 PDX when cultured under 3D *ex-vivo* conditions. However, consistent with our data from figure 5, the overall growth of these T-DM1-resistant PDX *ex-vivo* cultures was not inhibited by lapatinib treatment (Fig. 7D). In contrast, treatment of these *ex-vivo* cultures or tumor bearing mice with FIIN4, the covalent FGFR inhibitor, led to significant inhibition of tumor growth (Fig. 7b, 7d). Consistent with our previous studies, FIIN4 also demonstrated enhanced potency as compared to an identical concentration of its ATP competitive structural analogue, BGJ-398 (Fig. 7d). Taken together these data indicate that T-DM1 resistant tumors can be effectively targeted via covalent inhibition of FGFR kinase activity.

**Figure 7.**
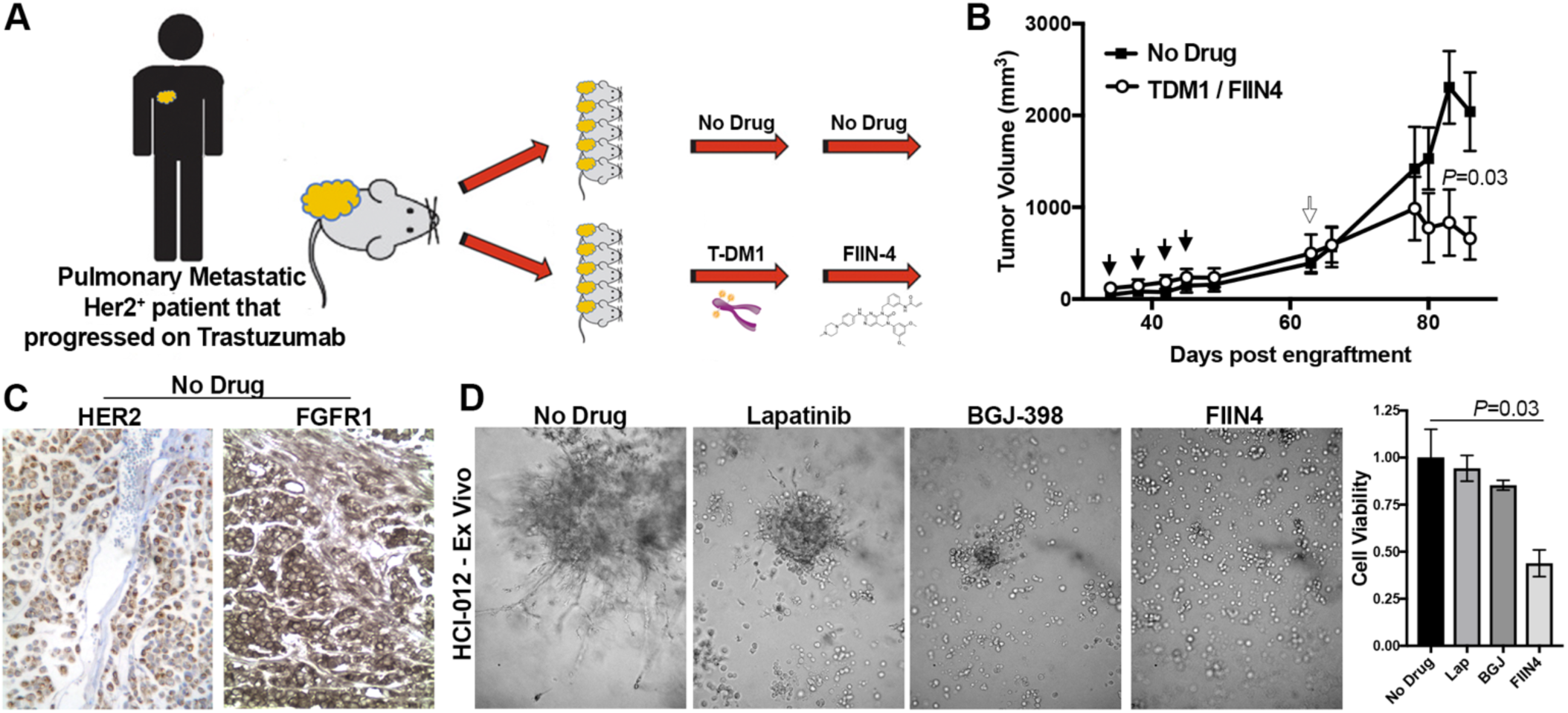
Trastuzumab-resistant patient derived xenografts are sensitive to covalent inhibition of FGFR. **A,** Schematic representation of the expansion protocol of HCI-012. Tumor bearing mice were split into two cohorts consisting of an untreated group and a group that was initially treated with T-DM1 (5 mg/kg) and then switched to FIIN4 (25 mg/kg/p.o.d). **B,** PDX tumor bearing mice were treated as described in panel A and tumor size was measured by digital caliper measurements at the indicated time points. Closed arrows indicate T-DM1 treatments, open arrow indicates initiation of FIIN4 treatment. Data are the mean ±SE of 5 mice per group resulting in the indicated p-value. **C,** Representative histological sections of untreated HCI-012 tumors stained with antibodies for HER2 and FGFR1. **D,** *Ex-vivo* HCI-012 tumor cells were grown for 20 days under 3D culture conditions in the presence or absence of the indicated compounds. Representative images are shown and cell viability was quantified by cell titer glow. Data are mean ±SD of triplicate wells treated with the indicated compounds.

## Discussion

Genomic amplification and high-level protein expression of HER2 cause constitutive oncogenic activity in a significant subset of breast cancer patients. These molecular events have precipitated robust diagnostics for trastuzumab-based therapeutics in these patients. Following disease progression on trastuzumab, patients can be treated with T-DM1, lapatinib or neratinib. In contrast to this current progression, our data and previous studies suggest that sequencing of these therapies could be improved through the first line use of kinase inhibitors followed by HER2 antibodies and T-DM1(31). Indeed, cells that acquired resistance to lapatinib still express HER2 and are still sensitive to T-DM1. In contrast, upon acquisition of resistance to T-DM1, HER2 can be diminished and these cells are therefore inherently resistant to ErbB kinase inhibitors. All of these therapies are predicated on the assumption that tumors and their corresponding metastases will remain addicted to HER2 or other ErbB receptors. However, increasing numbers of experimental and clinical studies indicate that following T-DM1 or trastuzumab treatment and metastasis, tumors can lose HER2 expression and become discordant (4,32,33)(34). Similar findings have been observed in B-cell lymphoma where expression of CD20 is diminished following treatment with Ritixumab (35). The mechanisms of antibody target diminution following therapeutic relapse and/or metastasis remain to be definitively determined. However, our findings using an ectopic HER2 overexpression system in immunodeficient mice suggest that acquisition of HER2 discordance is an active process that does not require immune-mediated cell clearance or endogenous transcriptional elements. Overall, the concept of HER2-initated tumors being capable of undergoing recurrence in a HER2 independent, EMT-driven fashion is supported by recent studies using doxycycline regulated models of HER2 expression (36,37).

Herein, we demonstrate that FGFR1 can act as a major driver of tumor recurrence following onset of HER2 discordance and acquisition of resistance to T-DM1 and other ErbB-targeted therapies. Previous studies from our lab and others suggest that FGFR can act as an bypass mechanism to facilitate resistance to ErbB kinase inhibitors (17,18,20,38). Furthermore, FGFR signaling has also been identified as a mechanism of resistance to endocrine therapies in breast and prostate cancer (19,39). Therefore, FGFR appears to constitute a critical node in acquisition of drug resistance. The reasons for this are potentially numerous, but a possible explanation is the inducible nature of FGFR expression. In particular, FGFR1, FGFR3 and FGF2 expression are dramatically upregulated during the processes of EMT (17,20). While the mechanisms of FGFR1 upregulation during EMT remain to be fully elucidated, this event presents a functional and targetable link between EMT and the acquisition of drug resistance.

FGFR signaling is clearly capable of acting as a bypass pathway during the acquisition of resistance to ErbB kinase inhibitors. However, our current data suggest that enhanced FGFR1 expression also plays a more active role in manifesting T-DM1 resistance by disrupting trastuzumab’s ability to bind to HER2. Determining the mechanisms by which FGFR prevents trastuzumab binding is currently under investigation in our laboratory. If FGFR physically disrupts antibody binding, than additional means besides kinase inhibition, may be required to degrade FGFR, reestablish trastuzumab binding, and prevent TDM1 persistence. In contrast, FGFR signaling has been shown to enhance the expression of ADAM10, a matrixmetaloproteinase capable of diminishing cellular binding of trastuzumab via its cleavage of the extracellular portion HER2 (40). Our data do not rule out this kind of indirect mechanism of reduced trastuzumab binding. Our data do clearly show that diminished binding of trastuzumab to cells can be manifest by induction of EMT or by direct overexpress of FGFR1. Our *in vitro* studies in figure 2 demonstrate that prior induction of EMT with TGF-β1 is required for eventual emergence of a TDM1-resistant subpopulation following prolonged drug treatment. These findings are in stark contrast to our previous studies with ErbB kinase inhibitors in which prior induction of EMT with TGF-β1 led to immediate resistant to both lapatinib and afatinib (20). Overall our data are consistent with a model in which cytokine-induced EMT upregulates FGFR1 which can immediately serve as a bypass pathway to overcome inhibition of ErbB receptor kinase activity. However, FGFR1-mediated diminishment of trastuzumab binding also allows for the survival of a subpopulation of cells in the face of T-DM1 treatment. This drug persistent subpopulation can then give rise to a proliferative, HER2 negative population that is fully resistant to T-DM1. This presents a unique combination of both subpopulation selection and phenotype plasticity, two processes typically thought to be mutually exclusive in drug resistance.

Our studies present covalent inhibition of FGFR as a potential approach for targeting T-DM1 resistance tumors. We developed FIIN4 as the first-in-class covalent inhibitor of FGFR, and an additional covalent FGFR inhibitor, TAS-120, is currently in phase 2 clinical trials in patients with advanced solid tumors (NCT02052778)(24). These studies and the studies herein demonstrate the enhanced efficacy of covalent kinase inhibition of FGFR as compared to first generation ATP competitive inhibitors. This may be a result of the structural stabilization of the inactivate confirmation that results upon covalent engagement of FGFR by FIIN4 (41). However, erdafitinib, an extremely potent competitive inhibitor of FGFR has recently been FDA approved. Therefore, clinical investigation of sequential or direct combinations of FGFR inhibitors with T-DM1 in the HER2^+^ setting is possible and clearly warranted.

## Acknowledgments

This research was supported in part by the American Cancer Society (RSG-CSM130259) to MKW and the National Institutes of Health (R01CA207751; R01CA232589; R21AA026675) to MKW and the UAB Breast SPORE Career Development Award to CDW. We also acknowledge the support of the Purdue Center for Cancer Research via its NIH NCI grant (P30CA023168). We kindly acknowledge the expertise of the personnel within the Purdue Center for Cancer Research Biological Evaluation Core. We also acknowledge the use of the facilities within the Bindley Bioscience Center, a core facility of the NIH-funded Indiana Clinical and Translational Sciences Institute.

## Supplementary Figures

**Suppl.Fig. S1.**
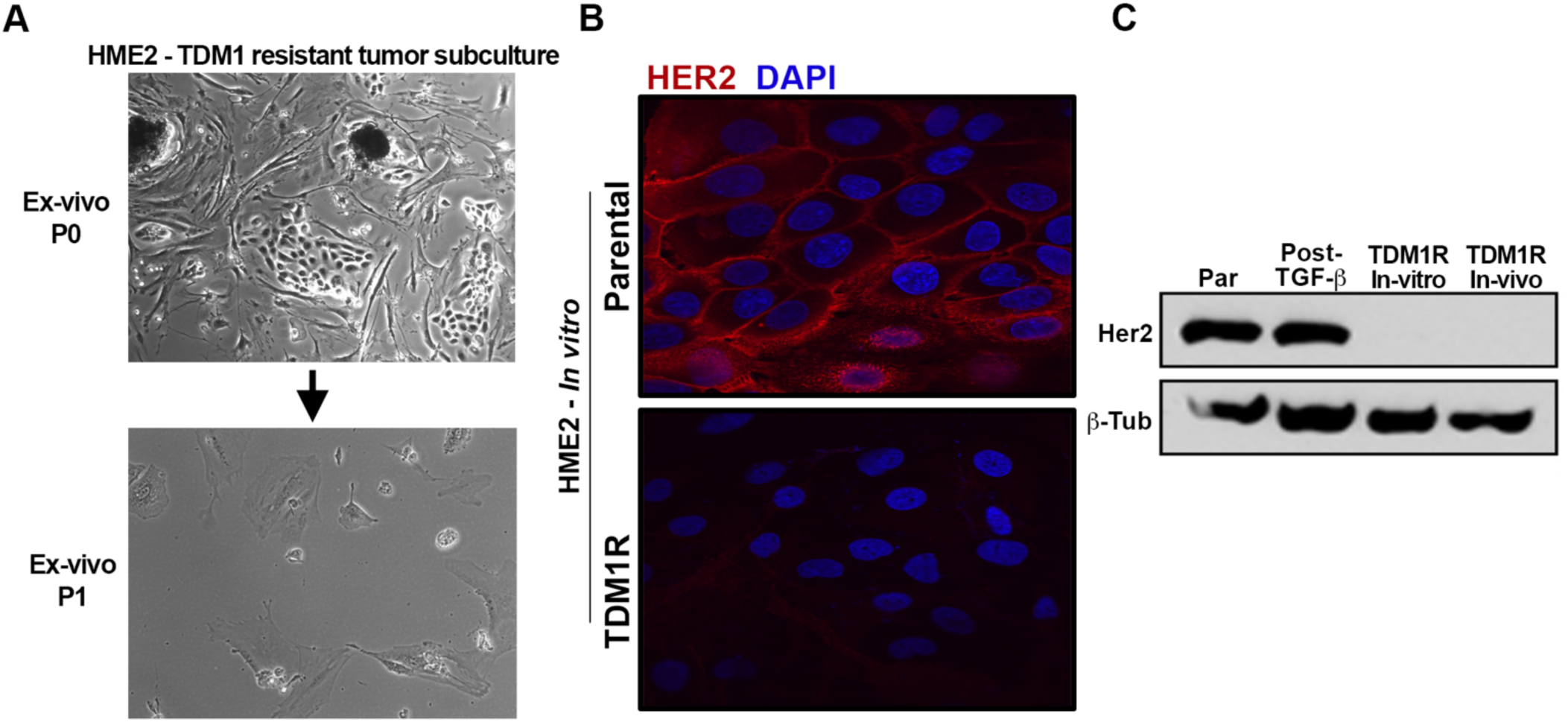
HER2 expression is diminished upon acquisition of resistantance to TDM1. **A,** *Ex-vivo* subculture of HME2 tumors that recurred after T-DM1-induced minimal residual disease as shown in Figure 1. These cells failed to thrive in culture. **B,** HME2 parental cells and their *in vitro*-derived T-DM1 resistant (TDM1R) counterparts were fixed, permeabilized and stained for HER2. These cells were counter stained with DAPI to visualize the nucleus. **C,** HME2 parental cells (Par), those treated and recovered from TGF-β1 (Post-TGF-β), those treated and recovered from TGF-β1 and subsequently selected for by continuous treatment with T-DM1 (TDM1R *In-vitro*), and primary culture from P3 HME2 tumors selected for resistance to T-DM1, as described in Figure 7a of the main text (TDM1R *In-vivo*), were assayed by immunoblot for expression of HER2 and β-tubulin (β-Tub) served as a loading control.

**Suppl.Fig. S2.**
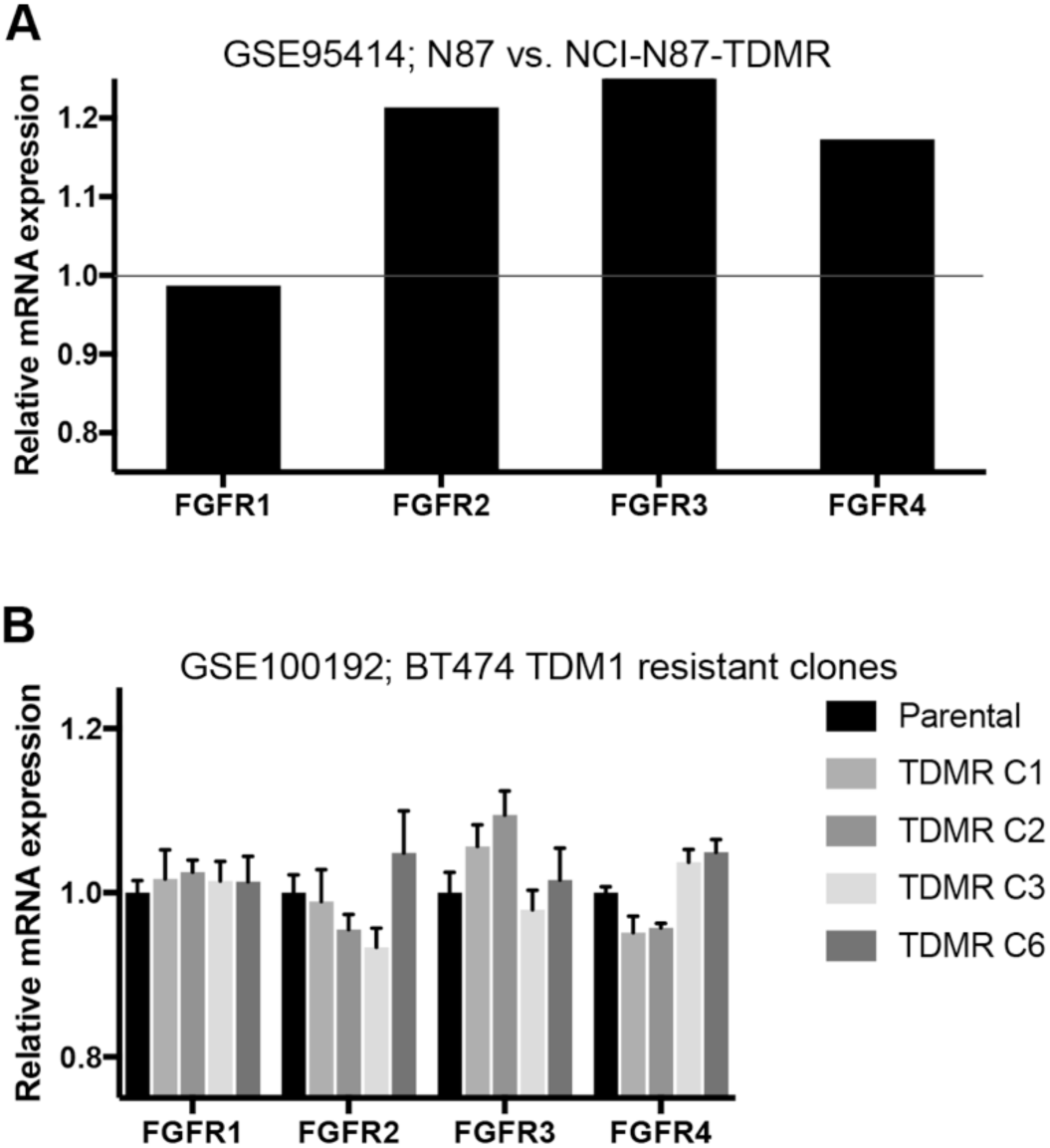
Enhanced FGFR expression upon acquisition of resistance to TDM1. **A,** Expression levels of FGFR1-4 in the NCI-N87 cells selected for resistant to TDM1. Data are extracted from a single RNA sequencing experiment and are normalized to the untreated control cells. **B,** Expression levels of FGFR1-4 in four different TDM1 resistant BT474 clones (C1-3 and C6). Data are mean ±SD of expression values from triplicate RNA sequencing experiments conducted for the parental cells and each TDM1 resistant clone.

**Suppl.Fig. S3.**
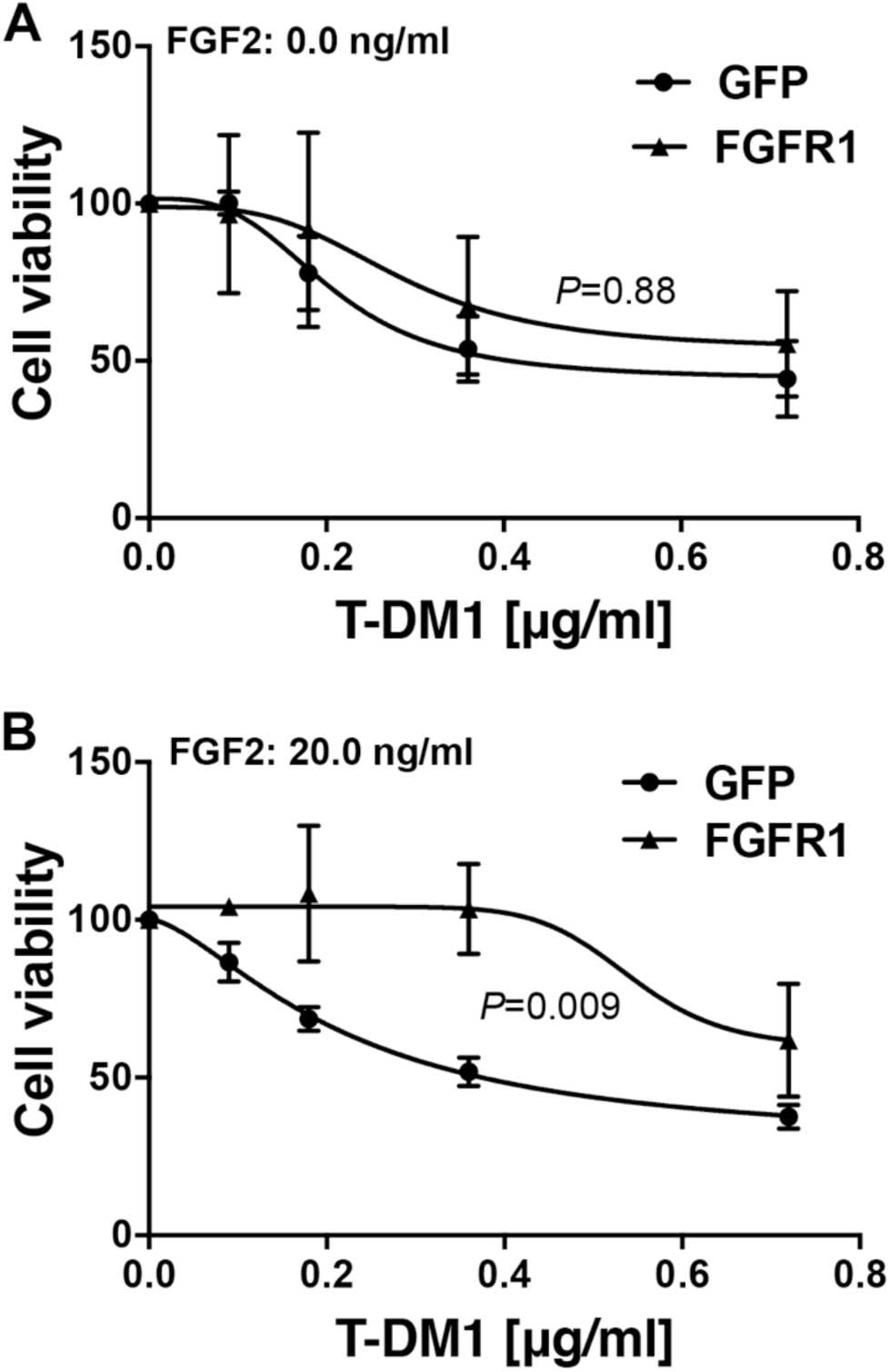
FGFR signaling is sufficient to diminish BT474 response to T-DM1. **A and B,** HER2 amplified BT474 cells expressing FGFR1 or GFP as a control were treated with the indicated concentrations of T-DM1 for 96 hours at which point cell viability was quantified. The dose response was done in the absence (**A**) and presence (**B**) of exogenous FGF2. Data are normalized to the untreated control cells and are the mean ±SEM of two independent experiments resulting in the indicated *P* value.

**Supplemental Table 1.**
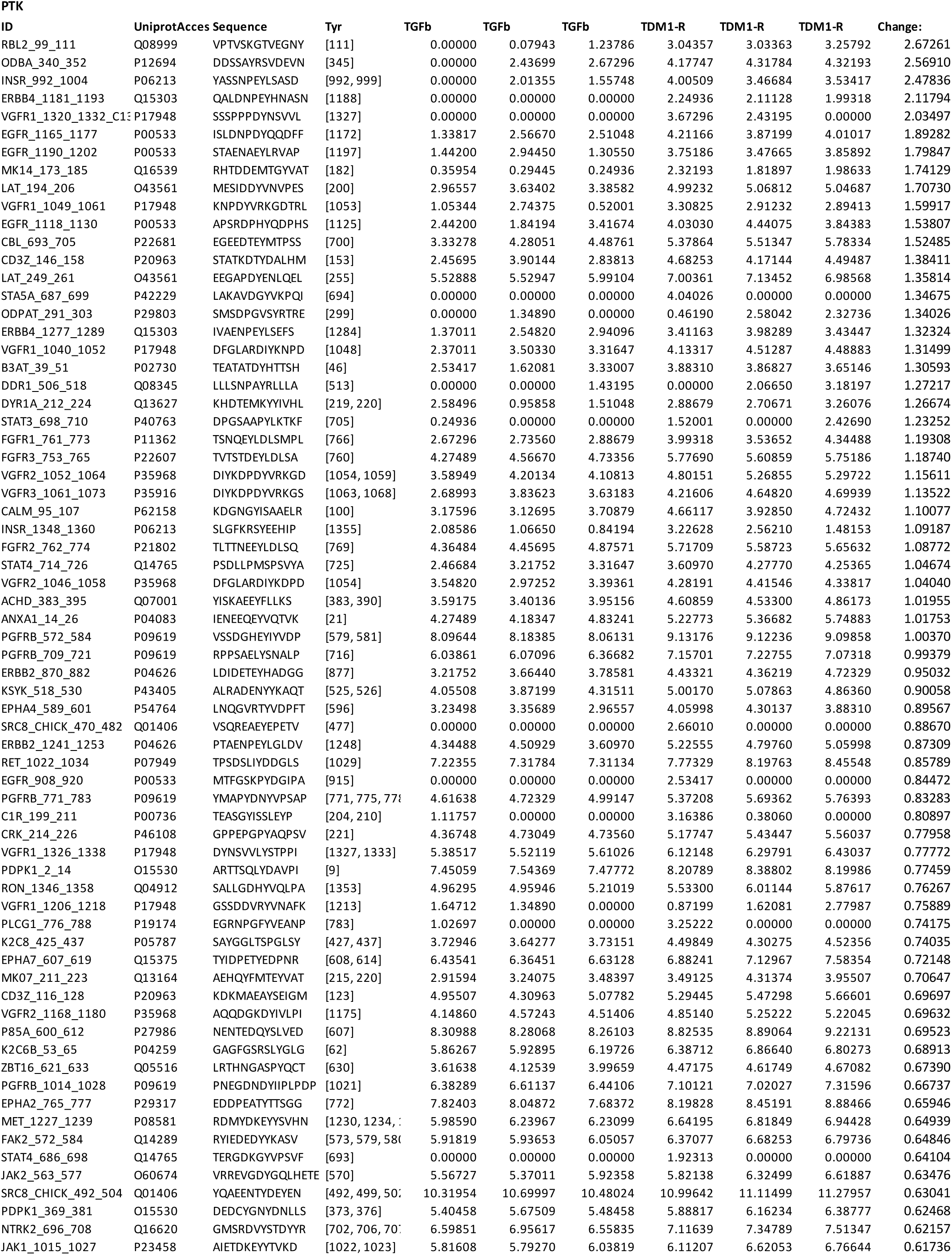
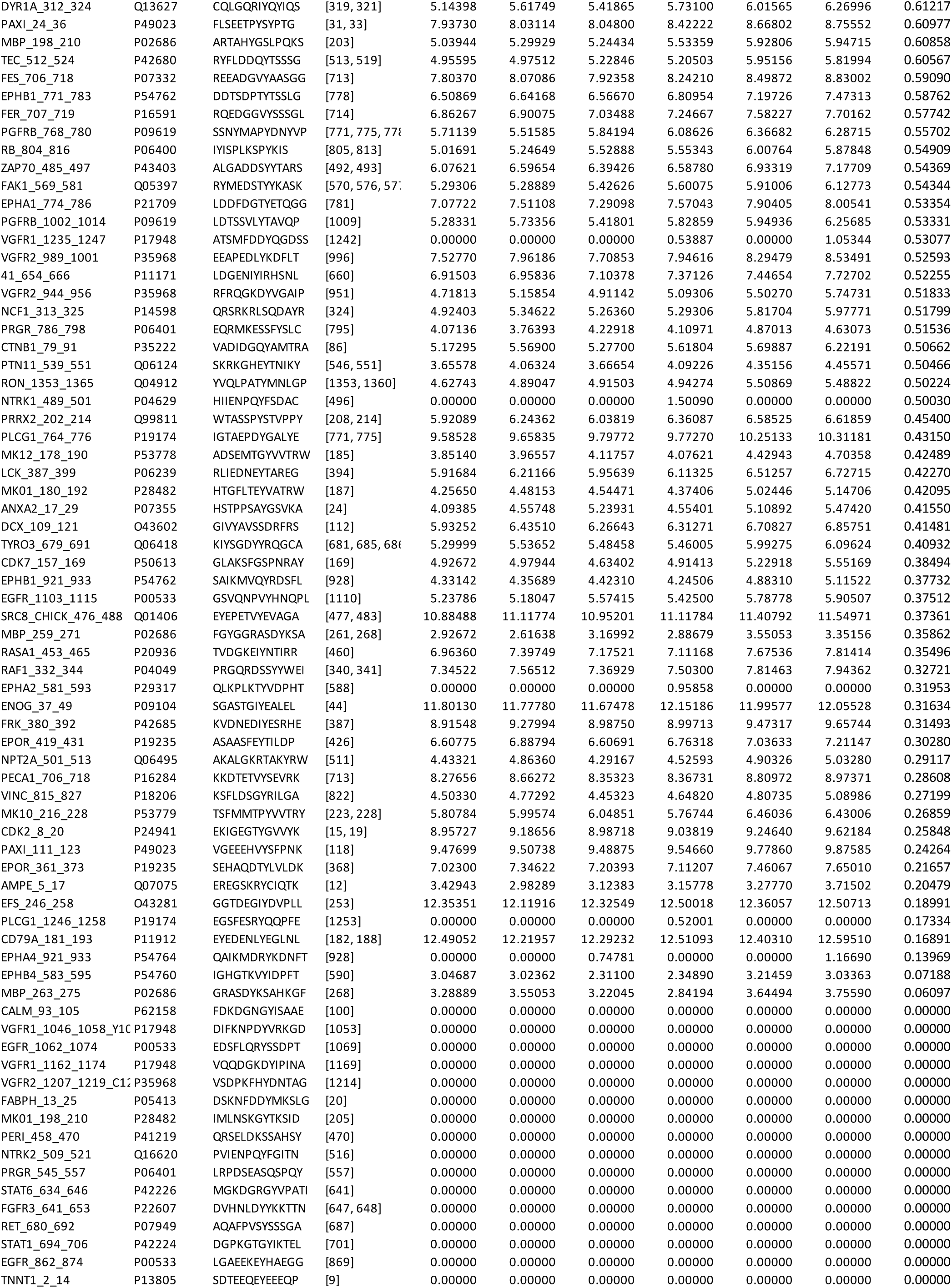
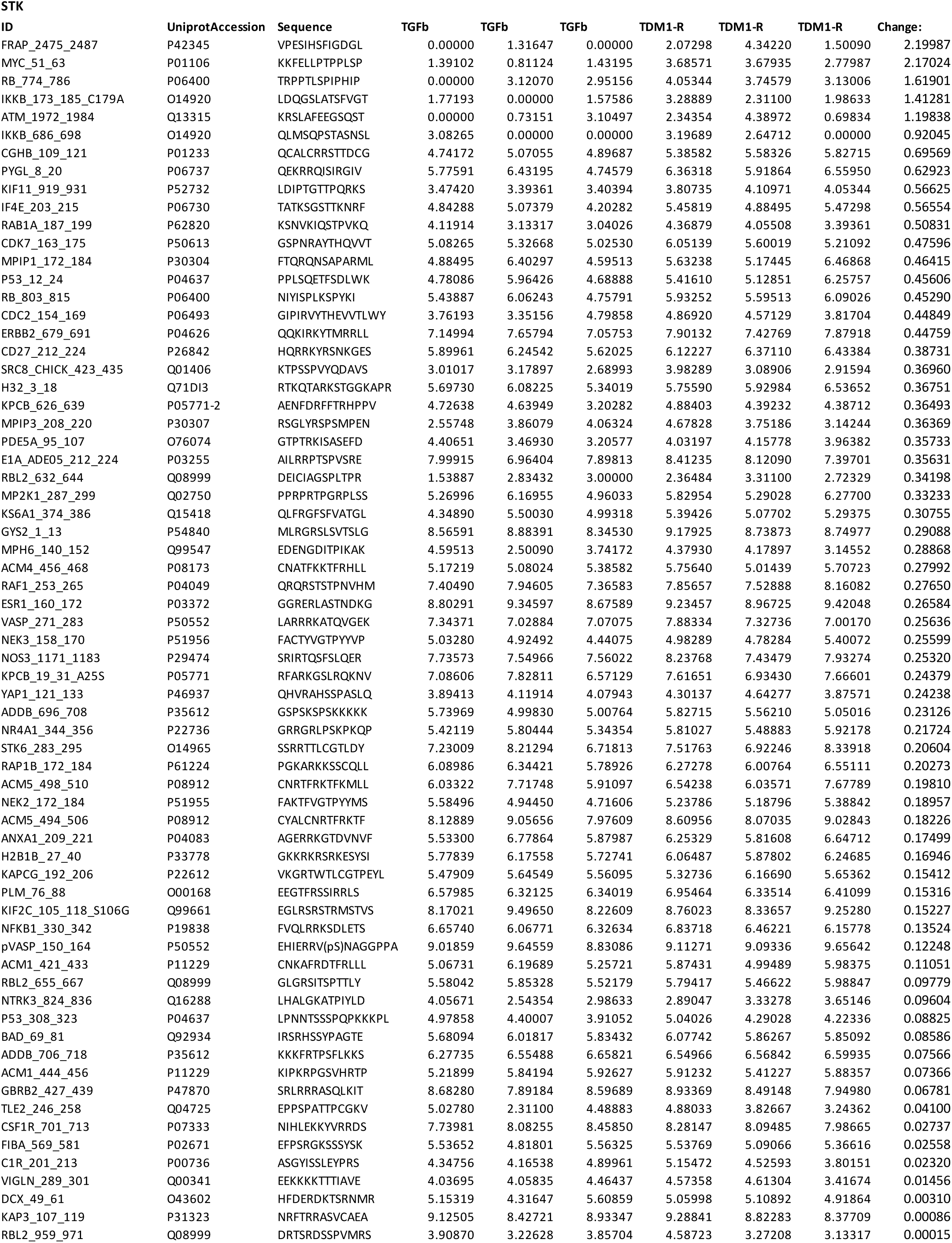
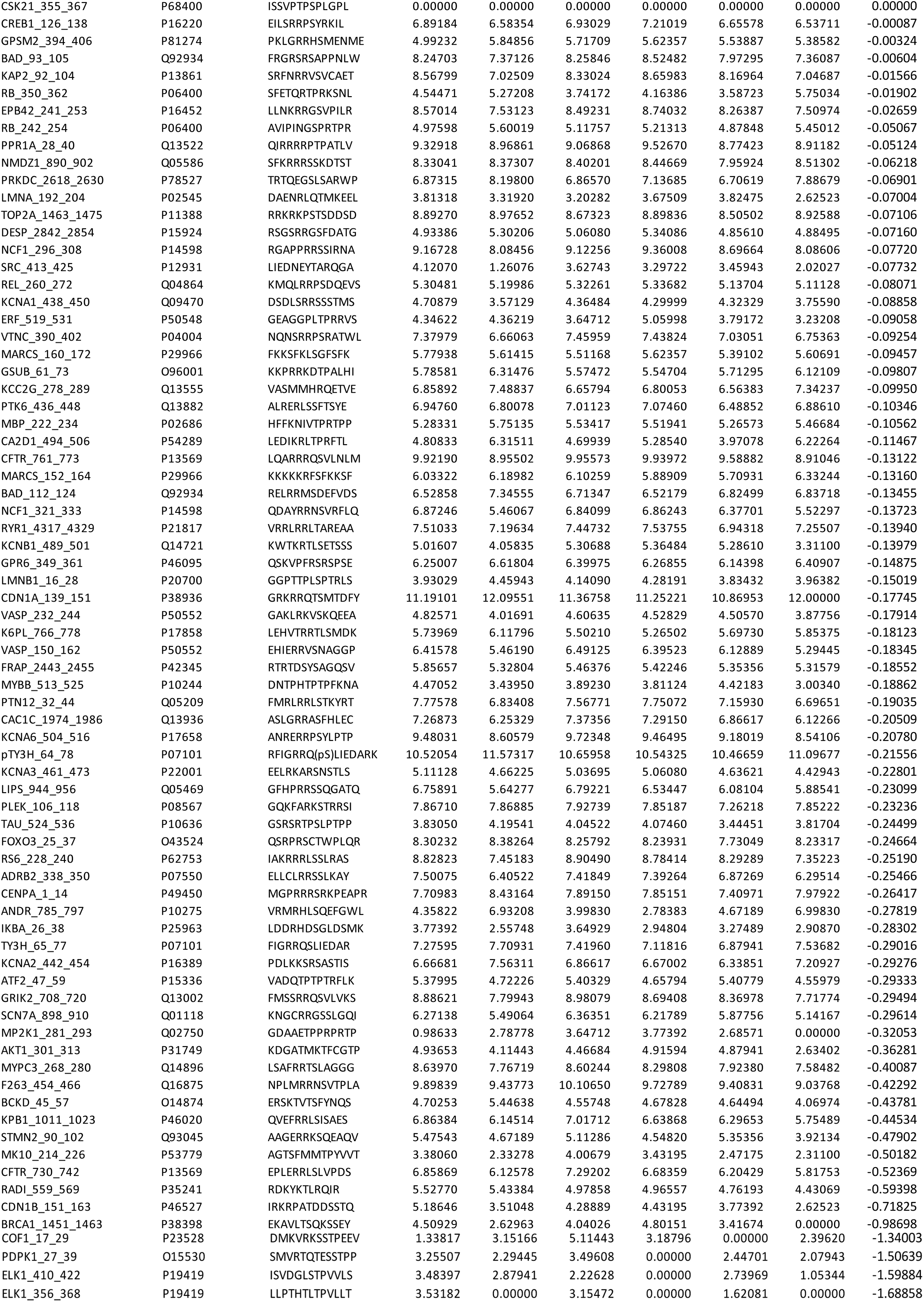
A list of differentially phosphorylated peptides from protein tyrosine kinases (PTK) and serine threonine kinases (STK). Triplicate values were obtained from TGF-β pretreated cells (TGFb) and the resulting T-DM1 resistant cells (TDM1-R). Peptides are ranked according to changes in phosphorylation levels between the two cell lines.

## References

1. Ado-Trastuzumab Emtansine Fails to Replace Standard of Care in First-Line Metastatic Breast Cancer - The ASCO Post [Internet]. [cited 2016 Jun 1]. Available from: http://www.ascopost.com/issues/july-10-2015/ado-trastuzumab-emtansine-fails-to-replace-standard-of-care-in-first-line-metastatic-breast-cancer/

2. Phase III MARIANNE Trial Results [Internet]. ASCO Annu. Meet. 2015 [cited 2016 Jun 1]. Available from: http://am.asco.org/phase-iii-marianne-trial-results

3. Barok M, Joensuu H, Isola J. Trastuzumab emtansine: mechanisms of action and drug resistance. Breast Cancer Res BCR. 2014;16:209.

4. Niikura N, Liu J, Hayashi N, Mittendorf EA, Gong Y, Palla SL, et al. Loss of Human Epidermal Growth Factor Receptor 2 (HER2) Expression in Metastatic Sites of HER2-Overexpressing Primary Breast Tumors. J Clin Oncol. 2011;JCO.2010.33.8889.

5. Choong L-Y, Lim S, Loh MC-S, Man X, Chen Y, Toy W, et al. Progressive loss of epidermal growth factor receptor in a subpopulation of breast cancers: implications in target-directed therapeutics. Mol Cancer Ther. 2007;6:2828–42.

6. Wendt MK, Tian M, Schiemann WP. Deconstructing the mechanisms and consequences of TGF-β-induced EMT during cancer progression. Cell Tissue Res. 2012;347:85–101.

7. Wendt MK, Smith JA, Schiemann WP. Transforming growth factor-β-induced epithelial-mesenchymal transition facilitates epidermal growth factor-dependent breast cancer progression. Oncogene. 2010;29:6485–98.

8. Wendt MK, Taylor MA, Schiemann BJ, Schiemann WP. Down-regulation of epithelial cadherin is required to initiate metastatic outgrowth of breast cancer. Mol Biol Cell. 2011;22:2423–35.

9. Wendt MK, Schiemann BJ, Parvani JG, Lee Y-H, Kang Y, Schiemann WP. TGF-β stimulates Pyk2 expression as part of an epithelial-mesenchymal transition program required for metastatic outgrowth of breast cancer. Oncogene. 2013;32:2005–15.

10. Kalluri R, Weinberg RA. The basics of epithelial-mesenchymal transition. J Clin Invest. 2009;119:1420–8.

11. Sharma SV, Lee DY, Li B, Quinlan MP, Takahashi F, Maheswaran S, et al. A chromatin-mediated reversible drug-tolerant state in cancer cell subpopulations. Cell. 2010;141:69–80.

12. Wendt MK, Taylor MA, Schiemann BJ, Sossey-Alaoui K, Schiemann WP. Fibroblast growth factor receptor splice variants are stable markers of oncogenic transforming growth factor β1 signaling in metastatic breast cancers. Breast Cancer Res BCR. 2014;16:R24.

13. Warzecha CC, Jiang P, Amirikian K, Dittmar KA, Lu H, Shen S, et al. An ESRP-regulated splicing programme is abrogated during the epithelial-mesenchymal transition. EMBO J. 2010;29:3286–300.

14. Madden SF, Clarke C, Gaule P, Aherne ST, O’Donovan N, Clynes M, et al. BreastMark: an integrated approach to mining publicly available transcriptomic datasets relating to breast cancer outcome. Breast Cancer Res BCR. 2013;15:R52.

15. Elbauomy Elsheikh S, Green AR, Lambros MBK, Turner NC, Grainge MJ, Powe D, et al. FGFR1 amplification in breast carcinomas: a chromogenic in situ hybridisation analysis. Breast Cancer Res BCR. 2007;9:R23.

16. Turner N, Pearson A, Sharpe R, Lambros M, Geyer F, Lopez-Garcia MA, et al. FGFR1 amplification drives endocrine therapy resistance and is a therapeutic target in breast cancer. Cancer Res. 2010;70:2085–94.

17. Raoof S, Mulford IJ, Frisco-Cabanos H, Nangia V, Timonina D, Labrot E, et al. Targeting FGFR overcomes EMT-mediated resistance in EGFR mutant non-small cell lung cancer. Oncogene. 2019;1.

18. Hanker AB, Garrett JT, Estrada MV, Moore PD, Ericsson PG, Koch JP, et al. HER2-Overexpressing Breast Cancers Amplify FGFR Signaling upon Acquisition of Resistance to Dual Therapeutic Blockade of HER2. Clin Cancer Res Off J Am Assoc Cancer Res. 2017;23:4323–34.

19. Mao P, Cohen O, Kowalski KJ, Kusiel JG, Buendia-Buendia JE, Exman P, et al. Acquired FGFR and FGF alterations confer resistance to estrogen receptor (ER) targeted therapy in ER+ metastatic breast cancer. bioRxiv. 2019;605436.

20. Brown WS, Akhand SS, Wendt MK, Brown WS, Salehin Akhand S, Wendt MK. FGFR signaling maintains a drug persistent cell population following epithelial-mesenchymal transition. Oncotarget. 2016;7:83424–36.

21. Brown, Wells, Li T, Smith, Andrew, Nathanael G, Wendt, Micahel. Covalent targeting of fibroblast growth factor receptor inhibits metastatic breast cancer. Mol Cancer Ther. In Press;

22. Dey JH, Bianchi F, Voshol J, Bonenfant D, Oakeley EJ, Hynes NE. Targeting fibroblast growth factor receptors blocks PI3K/AKT signaling, induces apoptosis, and impairs mammary tumor outgrowth and metastasis. Cancer Res. 2010;70:4151–62.

23. Sharpe R, Pearson A, Herrera-Abreu MT, Johnson D, Mackay A, Welti JC, et al. FGFR signaling promotes the growth of triple negative and basal-like breast cancer cell lines both in vitro and in vivo. Clin Cancer Res Off J Am Assoc Cancer Res. 2011;17:5275–86.

24. Brown WS, Tan L, Smith A, Gray NS, Wendt MK. Covalent Targeting of Fibroblast Growth Factor Receptor Inhibits Metastatic Breast Cancer. Mol Cancer Ther. 2016;15:2096–106.

25. Gilbert AN, Anderson JC, Duarte CW, Shevin RS, Langford CP, Singh R, et al. Combinatorial Drug Testing in 3D Microtumors Derived from GBM Patient-Derived Xenografts Reveals Cytotoxic Synergy in Pharmacokinomics-informed Pathway Interactions. Sci Rep. 2018;8:8412.

26. Shinde A, Hardy SD, Kim D, Akhand SS, Jolly MK, Wang W-H, et al. Spleen tyrosine kinase-mediated autophagy is required for epithelial-mesenchymal plasticity and metastasis in breast cancer. Cancer Res. 2019;

27. Mani SA, Guo W, Liao M-J, Eaton EN, Ayyanan A, Zhou AY, et al. The epithelial-mesenchymal transition generates cells with properties of stem cells. Cell. 2008;133:704–15.

28. Gámez-Pozo A, Pérez Carrión RM, Manso L, Crespo C, Mendiola C, López-Vacas R, et al. The Long-HER study: clinical and molecular analysis of patients with HER2+ advanced breast cancer who become longterm survivors with trastuzumab-based therapy. PloS One. 2014;9:e109611.

29. Rusnak DW, Lackey K, Affleck K, Wood ER, Alligood KJ, Rhodes N, et al. The Effects of the Novel, Reversible Epidermal Growth Factor Receptor/ErbB-2 Tyrosine Kinase Inhibitor, GW2016, on the Growth of Human Normal and Tumor-derived Cell Lines in Vitro and in Vivo. Mol Cancer Ther. 2001;1:85–94.

30. Hirsh V. Next-Generation Covalent Irreversible Kinase Inhibitors in NSCLC: Focus on Afatinib. Biodrugs. 2015;29:167–83.

31. Junttila TT, Li G, Parsons K, Phillips GL, Sliwkowski MX. Trastuzumab-DM1 (T-DM1) retains all the mechanisms of action of trastuzumab and efficiently inhibits growth of lapatinib insensitive breast cancer. Breast Cancer Res Treat. 2011;128:347–56.

32. Xiao C, Gong Y, Han EY, Gonzalez-Angulo AM, Sneige N. Stability of HER2-positive status in breast carcinoma: a comparison between primary and paired metastatic tumors with regard to the possible impact of intervening trastuzumab treatment. Ann Oncol Off J Eur Soc Med Oncol ESMO. 2011;22:1547–53.

33. Arihiro K, Oda M, Ogawa K, Tominaga K, Kaneko Y, Shimizu T, et al. Discordant HER2 status between primary breast carcinoma and recurrent/metastatic tumors using fluorescence in situ hybridization on cytological samples. Jpn J Clin Oncol. 2013;43:55–62.

34. Li G, Guo J, Shen B-Q, Yadav DB, Sliwkowski MX, Crocker LM, et al. Mechanisms of Acquired Resistance to Trastuzumab Emtansine in Breast Cancer Cells. Mol Cancer Ther. 2018;17:1441–53.

35. Duman BB, Sahin B, Ergin M, Guvenc B. Loss of CD20 antigen expression after rituximab therapy of CD20 positive B cell lymphoma (diffuse large B cell extranodal marginal zone lymphoma combination): a case report and review of the literature. Med Oncol Northwood Lond Engl. 2012;29:1223–6.

36. Abravanel DL, Belka GK, Pan T, Pant DK, Collins MA, Sterner CJ, et al. Notch promotes recurrence of dormant tumor cells following HER2/neu-targeted therapy. J Clin Invest. 2015;125:2484–96.

37. Mabe NW, Fox DB, Lupo R, Decker AE, Phelps SN, Thompson JW, et al. Epigenetic silencing of tumor suppressor Par-4 promotes chemoresistance in recurrent breast cancer. J Clin Invest. 2018;128:4413–28.

38. Azuma K, Kawahara A, Sonoda K, Nakashima K, Tashiro K, Watari K, et al. FGFR1 activation is an escape mechanism in human lung cancer cells resistant to afatinib, a pan-EGFR family kinase inhibitor. Oncotarget. 2014;5:5908–19.

39. Bluemn EG, Coleman IM, Lucas JM, Coleman RT, Hernandez-Lopez S, Tharakan R, et al. Androgen Receptor Pathway-Independent Prostate Cancer Is Sustained through FGF Signaling. Cancer Cell. 2017;32:474-489.e6.

40. Wei W, Liu W, Serra S, Asa SL, Ezzat S. The breast cancer susceptibility FGFR2 provides an alternate mode of HER2 activation. Oncogene. 2015;

41. Tan L, Wang J, Tanizaki J, Huang Z, Aref AR, Rusan M, et al. Development of covalent inhibitors that can overcome resistance to first-generation FGFR kinase inhibitors. Proc Natl Acad Sci U S A. 2014;111:E4869–4877.

